# Isotemporal Substitution Effects of 24-hour Movement Behaviors on Executive Function in Preschool Children

**DOI:** 10.1101/2025.09.17.676915

**Authors:** Ling Wang, LUO Wanhong, Liu Yong

## Abstract

**Objective:** This study aims to examine the relationship between 24-h movement behaviors (MBs) and executive function (EF) in preschool children.

**Methods:** A total of 266 preschool children participated in this study. Physical activity (PA) and sedentary behavior (SB) were measured using an accelerometer (ActiGraph GT3X-BT), sleep duration was assessed using sleep logs, and EF was evaluated using the Early Years Toolbox. Compositional data analysis was employed to explore associations among the three movement behaviors.

**Results:** (1) The relative distribution of 24-h MBs was significantly associated with inhibition, shifting, and updating (all *p* < 0.001), with a model explanatory power > 10%. The explanatory power was the highest for inhibition (16.3%). (2) After adjusting for other MBs, SB was negatively associated with inhibition (γ12 = –0.11, *p* < 0.001), shifting (γ12 = –1.45, *p* = 0.001), and updating (γ12 = –0.58, *p* = 0.004). In contrast, sleep (SP) was positively associated with shifting (γ11 = 1.60, *p* = 0.020). (3) When 15 min/day of SB was isotemporally substituted with SP, inhibition scores increased by 0.003, and shifting scores increased by 0.064; conversely, replacing SP with SB resulted in a significant decline.

**Conclusion:** The relative distribution of 24-h MBs significantly predicted EF in preschool children. Among MBs, SB and SP were most strongly associated with EF. Shifting SB toward SP may effectively improve EF performance, particularly shifting ability.

## 1 Introduction

Executive function (EF) is a foundational construct rooted in neuropsychology, first identified by observing patients who sustained frontal lobe injuries during wartime, manifesting pronounced impairments in self-control and behavioral execution, hence the coining of the term “executive function” (Posne & Rothbart, 2000; Shimamura & Jurica, 1994). Despite extensive study, the academic community has not reached consensus on a uniform definition of EF. The concept is often distinguished along two dimensions: in its broad sense, EF encompasses the coordinated orchestration of multiple cognitive processes during information processing, whereas in its narrow sense, it typically refers to inhibitory control (Li & Bai, 2005). EF, or cognitive control, is a conscious, top-down neurocognitive mechanism that encompasses three core components: cognitive flexibility, inhibitory control, and working memory. These components facilitate goal-directed regulation of thoughts, actions, and emotions (Miyake et al., 2000). This tripartite framework has achieved widespread acceptance; therefore, this study operationalizes EF in three sub-functions: cognitive flexibility (shifting), inhibitory control (inhibition), and working memory (updating). EF, a high-level cognitive capability, develops rapidly during early childhood (Diamond, 2016; Luan, Wang, & Lu, 2013).

Extensive evidence has underscored the lasting impact of early childhood EF on various later-life outcomes. Early EF development correlates strongly with school readiness (Graziano et al., 2016; Li & Liu, 2013) and academic performance (Nelson et al., 2017), as well as with adolescent behavioral patterns and broader adult outcomes, including health, socioeconomic status, and quality of life (Moffitt et al., 2011). EF demonstrates considerable plasticity, especially during early childhood (Diamond & Ling, 2016). Emerging research points to 24-h movement behaviors (MBs), including physical activity (PA), sedentary behavior (SB), and sleep, as potentially critical influences on EF development (Bezerra et al., 2020; McNeill et al., 2020). Furthermore, MBs have been linked to diverse cognitive and psychological outcomes (Kuzik et al., 2017, Kuzik et al., 2020; Chang & Wang, 2022).

However, most extant studies have focused on the effects of individual behaviors in isolation, such as sleep (Bernier et al., 2021; Xing et al., 2018; Xing et al., 2022), SB (Cui et al., 2021), and PA (Cook et al., 2019; McNeill et al., 2020; Qu et al., 2020), without adequately addressing how these behaviors interact or compose across an entire day. This fragmented approach limits our understanding of how integrated daily patterns of MB influence EF. Accordingly, there is a growing recognition of the need for a holistic perspective that considers the combined effects of PA, SB, and sleep to inform more effective intervention strategies.

Recent studies using different methods have addressed this gap. McNeill et al. (2020) examined adherence to 24-h movement guidelines and its relationship with EF and psychosocial health in preschoolers, although the results for cognitive shifting were mixed. A Canadian study found that meeting both sleep and PA guidelines was related to overall developmental benefits and response inhibition, although SB had no direct effect (Kuzik et al., 2022). Lau et al. (2024) employed compositional and isotemporal reallocation analyses to assess associations between MBs and EF, identifying significant linkages and demonstrating the practicality of reallocating time among behaviors to benefit EF outcomes. Additionally, a cross-sectional study of Chinese children revealed that meeting more of the 24-h movement guidelines, primarily via PA and sleep, has been positively associated with EF, with screen time having no significant effect (Zeng et al., 2022). A recent study found that reallocating time from sleep or SB to moderate-to-vigorous physical activity (MVPA) enhances cognitive flexibility, affirming the importance of time-use intensity in EF development (Lu et al., 2023).

Nevertheless, existing findings remain inconsistent, partly due to variations in measurement tools, participant characteristics, and analytical approaches (Chang, Wang, & Zhu, 2025). To address these gaps, this study sought to extend the current body of knowledge by systematically investigating the associations between 24-h MBs and objectively assessed EF subdomains in preschool children. This study aimed to quantify the potential effects of reallocating time among different behaviors on EF outcomes using compositional data analysis and isotemporal substitution modeling. The anticipated results may advance theoretical understanding and offer an empirical foundation for developing integrated interventions to promote EF in early childhood.

## 2 Participants and Methods

### 2.1 Participants

The recruitment for this study was conducted from from September 9, 2024, to January 24, 2025. A total of 371 preschool children were initially recruited through convenience sampling from one public kindergarten and two private kindergartens in Changsha City. Subsequently, Following this, 319 parents provided written informed consent for their own and their children’s participation in the study. After completing the 24-hour movement behavior assessment and questionnaire survey, valid data were obtained from 266 participants. The demographic characteristics of the final sample are summarized in Table 1. All study procedures, including participant recruitment, informed consent, testing protocols, and safety contingency plans, were reviewed and approved by the Human Research Ethics Committee of East China Normal University (Approval No. HR 342-2024).

**Table 1.**
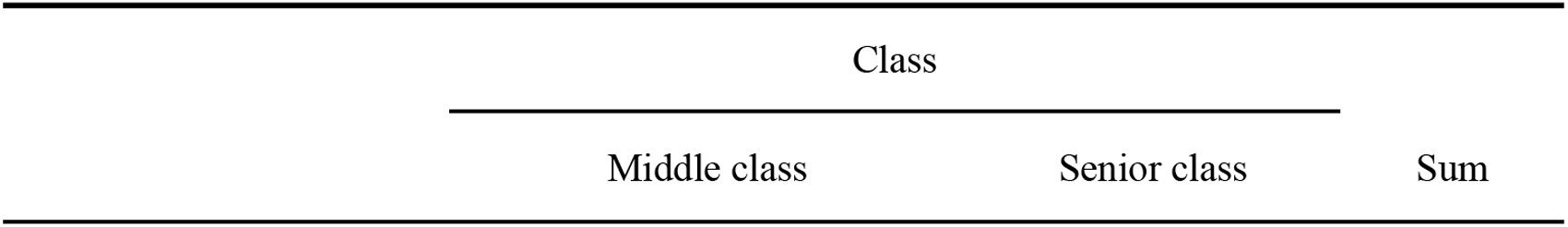
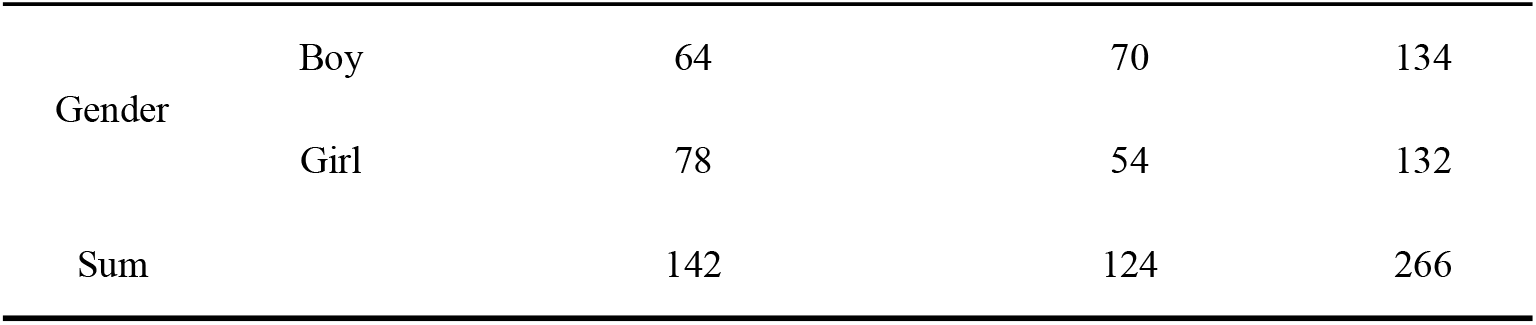
Demographic characteristics of the participants.

| Class |  |  |
| --- | --- | --- |
| Middle class | Senior class | Sum |

|  |  |  |  |  |
| --- | --- | --- | --- | --- |
| Gender | Boy | 64 | 70 | 134 |
|  | Girl | 78 | 54 | 132 |
| Sum |  | 142 | 124 | 266 |

### 2.2 Methods

#### 2.2.1 24-h MB Monitoring

A triaxial accelerometer (ActiGraph GT3X-BT, Pensacola, FL, United States) was used to obtain objective measures of PA and SB. This device provides estimates of light-intensity physical activity (LPA) and MVPA. Children were asked to wear the accelerometer on the right hip at the level of the iliac crest for seven consecutive days (five weekdays and two weekend days). Before data collection, parents and teachers received detailed instructions, highlighting that the device should be worn continuously, except during sleep or water-based activities.

Raw data were processed using ActiLife software. Non-wear time was determined using the Choi algorithm (Choi, Liu, Matthews, & Buchowski, 2011), and data were considered valid if children had at least three days of recording, including two weekdays and one weekend day, with a minimum daily wear time of 480 min. The epoch length was set as 15 s. Activity intensity cutoff points followed the thresholds proposed by Chang & Wang (2021): SB = 0–116 counts/15 s, LPA = 117–551 counts/15 s, MPA = 552–997 counts/15 s, and VPA ≥ 998 counts/15 s. These criteria have been validated in Chinese preschoolers aged three to six years.

Sleep duration was determined using an online collaborative sleep log jointly completed by the parents, head teachers, and classroom teachers. The log captured the children’s bedtimes, wake-up times, and nap onset and offset. Adapted from Chang and Wang (2023), this log displayed strong agreement with ActiGraph wGT3X-BT sleep estimates derived from the Tudor-Locke algorithm (r = 0.972, *p* < 0.001).

#### 2.2.2 EF Assessment

EF was assessed using the Early Years Toolbox (EYT) on an iPad. EYT is a validated and user-friendly tool that supports offline multi-device use and employs a gamified design to enhance children’s engagement (Howard & Melhuish, 2017). Assessments were conducted in a quiet, well-lit room, with each child accompanied by a trained assessor who provided standardized instructions.

EYT included three core tasks. (1) Card sorting: This task evaluates cognitive flexibility. Children were required to sort cards based on color or shape, with the sorting rule changing partway throughout the task. The score is based on the number of correct responses following the rule switches. (2) Go/No-Go: This task measures inhibitory control. Children were instructed to respond (“Go”) when a fish appeared and withhold response (“No-Go”) when a shark appeared. The probabilities of the fish and shark stimuli were 80% and 20%, respectively. The inhibition score was calculated as the product of the accuracy in Go and No-Go trials. (3) Mr. Ant: This task assesses working memory. Children had to remember the locations of the stickers on an ant’s body and identify them after a delay. The task included eight difficulty levels, with points awarded for each completed level. The task was terminated after three consecutive failures. All three tasks demonstrated good validity and were significantly correlated with the NIH Toolbox measures of working memory (r = 0.46), inhibition (r = 0.40), and cognitive flexibility (r = 0.45; all *p* < 0.001) (Howard & Melhuish, 2017).

#### 2.2.3 Covariates

Covariates included gender, age, and body mass index (BMI) of children. Gender and age information were obtained from class rosters. Height and weight were measured by kindergarten staff using standardized equipment (Jianmin brand) one week before testing. BMI was calculated using the following formula: BMI = weight (kg)/height (m2).

### 2.3 Data Analysis

Data processing and statistical analyses were conducted as follows. First, independent-sample t-tests and one-way analysis of variance were used to examine differences in EF and its subcomponents across demographic variables. Second, compositional data analysis (CoDA) was applied to describe the composition of 24-h MBs, including measures of central tendency (compositional mean) and dispersion (log-ratio variance). Third, MB data were transformed using the isometric log-ratio (ILR) transformation to investigate the associations between different MBs and EF. Regression models were established as follows: E(Y|Z) = β_0_ + β_1_z_1_ + β_2_z_2_ + β_3_z_3_ + … + β_(_d-1_)_z_(_d-1_)_ + Covariates; where Y represents EF or its subcomponents, d is the number of MB components, zi are the ILR-transformed variables, and βi are the corresponding regression coefficients. The covariates included gender, age, and BMI. Finally, R software (version 4.3.1) was used to analyze the isometric substitution effects of reallocating time between different MBs on EF, following the approach proposed by Dumuid et al. (2018, 2019).

## 3 Results

### 3.1 Basic Characteristics of EF in Preschool Children

Girls exhibited significantly higher inhibitory control than boys (t = –3.43, *p* = 0.001), with no significant differences in cognitive flexibility (t = 0.78, *p* = 0.434) or working memory (t = –0.85, *p* = 0.395).

Regarding age, preschool children aged three–four years performed significantly lower than those aged five–six years across all three EF dimensions, inhibitory control (t = –2.43, *p* = 0.016), cognitive flexibility (t = –3.05, *p* = 0.003), and working memory (t = –3.19, *p* = 0.002), indicating that EF improves with age.

Based on the World Health Organization (WHO) definition of overweight in children under five years (weight-for-height > 2 standard deviations above the WHO child growth standard median (https://www.who.int/zh/news-room/fact-sheets/detail/obesity-and-overweight), the prevalence of overweight in this sample was 10.5% (n = 28), higher than the 6.8% reported for children under six years in the Report on Nutrition and Chronic Disease Status of Chinese Residents (2020). However, no significant differences were observed in the overall EF or its subcomponents between the overweight and non-overweight children (Table 2).

**Table 2.** EF status of preschool children.

|  |  | N | Inhibitory control | Cognitive flexibility | Working memory |
| --- | --- | --- | --- | --- | --- |
| Gender | Boy | 13<br>4 | 0.85 ± 0.13** | 10.39 ± 1.40 | 2.48 ± 0.76 |
|  | Girl | 13<br>2 | 0.89 ± 0.87** | 10.24 ± 1.76 | 2.56 ± 0.70 |
| Age | 3–4 | 13<br>4 | 0.85 ± 0.13<sup>#</sup> | 10.03 ± 1.76<sup>##</sup> | 2.38 ± 0.78<sup>##</sup> |
|  | 5–6 | 13<br>2 | 0.89 ± 0.09<sup>#</sup> | 10.61 ± 1.33<sup>##</sup> | 2.66 ± 0.65<sup>##</sup> |
| BMI | Overweight | 28 | 0.86 ± 0.08 | 10.71 ± 1.12 | 2.43 ± 0.80 |
|  | Non-overweight | 23<br>8 | 0.87 ± 0.12 | 10.27 ± 1.63 | 2.53 ± 0.72 |
|  | Total | 26<br>6 | 0.87 ± 0.11 | 10.32 ± 1.58 | 2.52 ± 0.73 |

Note: \* indicates a significant difference between children of different genders. # indicates a significant difference between children of different age groups. \* and # represent $p < 0.05$ ; \*\* and ## represent $p < 0.01$ ; \*\*\* and ### represent $p < 0.001$ . BMI is divided into two parts according to WHO standards: overweight and non-overweight.

### 3.2 Characteristics of 24-h MBs in Preschool Children

The average composition of 24-h MBs among preschool children is presented in Table 3. Sleep accounted for 42.11% of the day, SB for 44.10%, LPA for 10.84%, and MVPA for 2.94%.

**Table 3.**
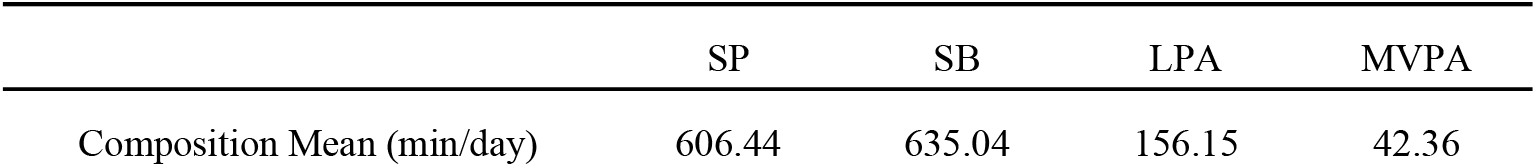
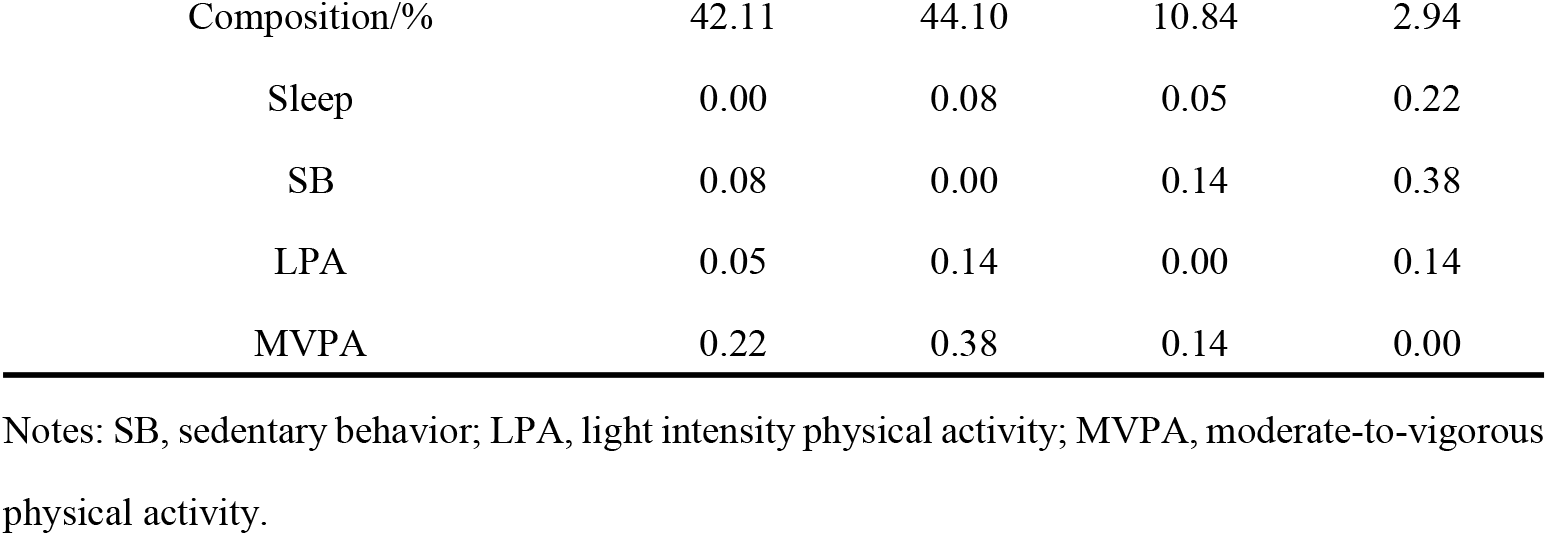
Compositional descriptive statistics of 24-h MBs: central tendency and dispersion measures.

The variation matrix analysis indicated that the log-ratio variance between SB and MVPA was the highest (0.38), suggesting a relatively high variability and low dependency between these two behaviors. Changes in MVPA time were more likely to be compensated by changes in LPA (log-ratio variance = 0.14). However, alterations in sleep time were likely replaced by SB (log-ratio variance = 0.08) and LPA (log-ratio variance = 0.05).

### 3.3 Associations Between 24-h MBs and EF

The associations between 24-h MBs and EF are presented in Table 4. The relative composition of 24-h MBs was significantly associated with inhibition, cognitive flexibility, and working memory (all *p* < 0.001), with model R^2^ values > 10% for each EF dimension. The highest model explanatory power was observed for inhibitory control (R^2^ = 16.3%). After controlling for other MBs, SB was negatively associated with inhibitory control (γ_1_^2^ = −0.11, *p* < 0.001), cognitive flexibility (γ_1_^2^ = −1.45, *p* = 0.001), and working memory (γ_1_^2^ = −0.58, *p* = 0.004). In contrast, sleep was positively associated with cognitive flexibility (γ_1_^1^ = 1.60, *p* = 0.020; Table 4).

**Table 4.**
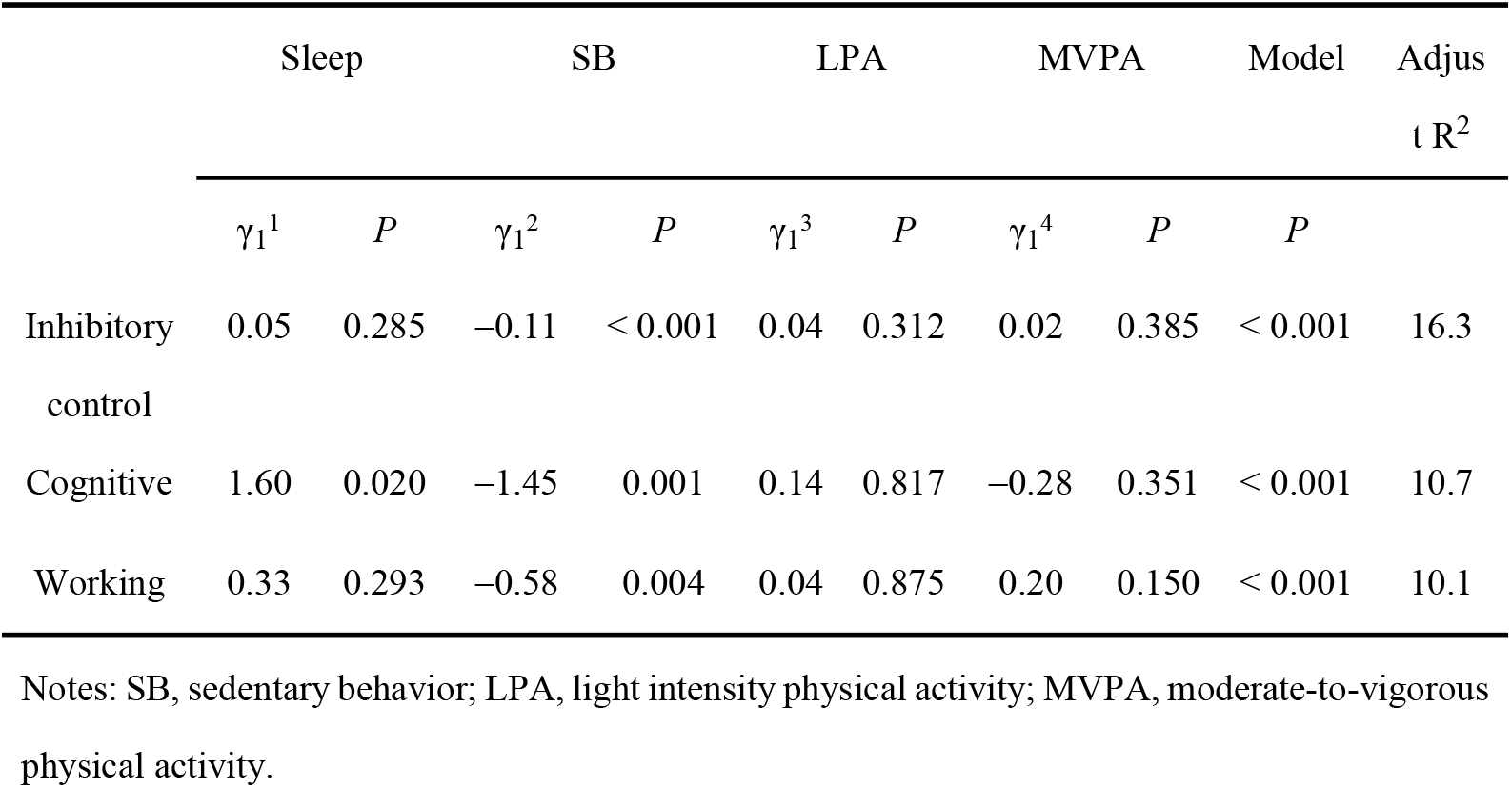
Associations between 24-h MBs and EF.

|  | Sleep |  | SB |  | LPA |  | MVPA |  | Model | Adjusted R<sup>2</sup> |
| --- | --- | --- | --- | --- | --- | --- | --- | --- | --- | --- |
| | $\gamma_1^1$ | *P* | $\gamma_1^2$ | *P* | $\gamma_1^3$ | *P* | $\gamma_1^4$ | *P* | *P* | |
| Inhibitory control | 0.05 | 0.285 | −0.11 | < 0.001 | 0.04 | 0.312 | 0.02 | 0.385 | < 0.001 | 16.3 |
| Cognitive | 1.60 | 0.020 | −1.45 | 0.001 | 0.14 | 0.817 | −0.28 | 0.351 | < 0.001 | 10.7 |
| Working | 0.33 | 0.293 | −0.58 | 0.004 | 0.04 | 0.875 | 0.20 | 0.150 | < 0.001 | 10.1 |

Notes: SB, sedentary behavior; LPA, light intensity physical activity; MVPA, moderate-to-vigorous physical activity.

### 3.4 Changes in EF Following 15-min/day Isotemporal Substitution of 24-h MBs

Following previous studies (Bezerra et al., 2020; Chang & Wang, 2022), an isotemporal substitution of 15 min/day was used to examine the effect of reallocating time between MBs on EF. The results indicated that replacing SB with sleep increased the EF scores of preschool children: inhibition increased by 0.003, and cognitive flexibility increased by 0.064. Conversely, reallocating time in the opposite direction resulted in significant decreases in the EF scores (Table 5).

**Table 5.** Effects of 15-min daily time reallocation between 24-h MBs on EF in preschool children.

| +15min | −15min | Inhibitory control | Cognitive flexibility | Working memory |
| --- | --- | --- | --- | --- |
| Sleep | SB | 0.003* | 0.064* | 0.019 |
| Sleep | LPA | −0.003 | 0.022 | 0.003 |
| Sleep | MVPA | −0.006 | 0.141 | −0.07 |
| SB | LPA | −0.006 | −0.041 | −0.015 |
| SB | MVPA | −0.009 | 0.078 | −0.088 |
| SB | Sleep | −0.003* | −0.064* | −0.019 |
| LPA | SB | 0.005 | 0.041 | 0.015 |
| LPA | MVPA | −0.004 | 0.118 | −0.073 |
| LPA | Sleep | 0.002 | −0.024 | −0.004 |
| MVPA | SB | 0.007 | −0.044 | 0.065 |
| MVPA | LPA | 0.001 | −0.086 | 0.049 |
| MVPA | Sleep | 0.004 | −0.109 | 0.046 |

Notes: SB, sedentary behavior; LPA, light intensity physical activity; MVPA, moderate-to-vigorous physical activity.

## 4. Discussion

This study examined the association between 24-h MBs and objectively measured EF in Chinese preschool children. Compared to prior fragmented studies focusing on individual behaviors, these findings highlight positive associations between sleep and EF, negative associations between SB and EF, and the effects of isotemporal substitution between these behaviors on EF.

EYT, developed in Australia for children aged three to five years, has been increasingly recognized for EF assessment in early childhood research (Howard & Melhuish, 2017). However, results from studies using EYT vary widely, with some reporting significant positive associations between MVPA and inhibitory control (Bezerra et al., 2020; Chang & Wang, 2020), while others found that vigorous physical activity (VPA) was positively related to cognitive flexibility (McNeill et al., 2020; Qu et al., 2020), and some reported no significant association between PA and inhibition control or cognitive flexibility, with even negative associations with working memory (Cook et al., 2019). These discrepancies suggest that the relationship between PA and EF may depend on the specific activity types and contextual factors.

To address these inconsistencies, some scholars have applied multivariate modeling approaches to systematically investigate the relationship between PA intensity spectra and EF in three-to five-year-old children. Analyses using uniaxial PA intensity data revealed that PA was significantly associated with inhibitory control (R^2^ = 1.48%), but not with working memory or cognitive flexibility. Specifically, the time spent in the 0–99 cpm and ≥ 8500 cpm intensity ranges was positively correlated with inhibitory control, whereas the time spent in the 100–3499 cpm range was negatively correlated. When the triaxial PA data were used, no significant association with EF was observed. Further stratified analyses revealed significant associations between PA and inhibitory control in girls (R^2^ = 3.12%) and in older children aged 4.6–6.5 years (R^2^ = 3.33%), although the directions of these associations varied across intensity ranges (Vabø, Aadland, Howard, & Aadland, 2022). These association patterns were difficult to interpret because SB and VPA were positively associated with EF in the same model, whereas LPA and MPA were negatively associated.

Thus, longitudinal studies may provide more reliable evidence. McNeill et al. (2020) reported that baseline VPA positively predicted cognitive flexibility after one year (b = 0.245; 95% CI: 0.006–0.485; *p* = 0.045), whereas MVPA exhibited a near-significant association (b = 0.119; 95% CI: –0.0001 to –0.239; *p* = 0.051). Children who did not participate in organized physical activities at baseline exhibited better inhibitory control after 12 months than those who did (Mdiff = 0.06; 95% CI: 0.00– 0.13; *p* = 0.046). Cross-sectional analyses further indicated that children meeting the 24-h movement guidelines performed better in verbal working memory (*p* = 0.026) and cognitive flexibility (*p* = 0.034), with stronger effects observed for those meeting two or three guideline components. Longitudinally, children who met the 24-h guidelines at baseline exhibited superior cognitive flexibility after 12 months compared to those who did not (*p* < 0.002) (McNeill et al., 2020).

Although the mechanisms linking PA and EF remain unclear, most studies suggest that PA intensity is positively associated with EF in preschool children. However, a significant association between PA and EF was not observed in this study. Several factors could explain this discrepancy. First, this study applied CoDA, treating 24-h MBs as a whole and accounting for the interdependence among behavioral components. Compared to traditional independent-variable approaches, CoDA provides a more comprehensive view of behavioral co-variation, possibly leading to different association directions. Second, the daily PA level of the children in this study was relatively low (∼42 min/day), potentially insufficient to reach the intensity threshold needed to significantly impact EF. Third, although isotemporal substitution can reveal the direction and pathway of behavior-time replacement effects on EF, it cannot fully characterize the dose-response relationship or determine the intensity-specific thresholds for PA effects (Wang et al., 2022). Nonetheless, the overall CoDA model indicated significant associations between 24-h time-use composition and all three EF dimensions, highlighting the importance of developing more refined statistical models to explore the complex mechanisms underlying PA and EF.

This study identified sleep and SB as key behaviors that are closely linked to EF in preschool children. Previous systematic reviews reported a weak positive correlation between sleep and EF (Reynaud et al., 2018). These findings revealed that sleep was positively associated with cognitive flexibility after controlling for other behaviors, and substituting 15 min/day of sleep with SB significantly improved inhibitory control and cognitive flexibility, supporting prior evidence. Regarding SB, existing studies report inconsistent findings; a review of 16 studies indicated that seven reported negative associations with EF and two reported positive associations (Li et al., 2022). In this study, SB was significantly negatively associated with all three EF dimensions, consistent with most previous research. From a neurobiological perspective, SB may impair white matter development, which is crucial for EF (Hutton et al., 2020). Given that myelination and neural signal conduction efficiency are influenced by environmental factors (Forbes & Gallo, 2017), prolonged SB may negatively impact EF development.

SB effects should be interpreted cautiously. Different types of sedentary activities may have divergent impacts on EF; cognitively stimulating activities (for instance, puzzles or problem-solving games) may enhance EF (Carson et al., 2015), whereas passive screen-based activities may be detrimental (Li et al., 2022). Neuroimaging studies have demonstrated that excessive screen time is associated with reduced frontal gray matter volume and decreased white matter anisotropy (Hutton et al., 2020; Zavala-Crichton et al., 2020), and the frontal lobe is a key brain region for cognitive flexibility (Cui, Li, & Dong, 2022). Overall, research on the neural mechanisms linking SB, especially cognitive flexibility and EF, remains limited and warrants further investigation (Cui et al., 2022; Li et al., 2022; Cui et al., 2021).

## 5 Implication and limitation

This study has several limitations. First, sleep duration was measured subjectively rather than objectively due to the inconvenience of wearing accelerometers overnight. This approach may have introduced some bias in the proportional estimates of 24-h MBs. Second, this study employed a cross-sectional design, which limits the ability to infer causal relationships. Third, EYT is relatively time-consuming, and the associated cognitive load may compromise assessment accuracy. A pilot study indicated that evaluating the three core EF dimensions required approximately 45 min per child, occasionally resulting in dropout due to fatigue or inattention (Bezerra et al., 2020). To address this issue, this study implemented several strategies, including scheduling assessments during children’s optimal activity periods, such as morning sessions, incorporating short breaks between tasks, and using trained assessors to maintain engagement. These measures helped reduce fatigue effects and improve data completeness and reliability.

## 6 Conclusion

The relative distribution of 24-h MBs significantly predicted EF in preschool children. Among these behaviors, SB and sleep exhibited particularly strong associations with all EF dimensions. Appropriately reducing SB while correspondingly increasing SP can effectively enhance EF, especially cognitive flexibility. These findings underscore the importance of optimizing the overall 24-h behavioral composition in practical interventions, highlighting that substituting sleep for sedentary time may promote the healthy development of EF in early childhood.

## Acknowledgment

Thank you for the strong support and assistance of the Three Education Series Kindergartens and Ronsheng Huayu City Kindergarten.

## Data availability

The datasets generated and/or analyzed during the current study are not publicly available according to the China legislation, as access to individual-level data is restricted only to individuals named in the study permit. The study protocol is available upon request from the corresponding author.

## Author information

Authors and Affiliations

LUO Wanhong, College of Physical Education, Hunan First Normal University Changsha, Hunan, PR China.

WANG Ling, College of Sports Science, Changsha Normal University, Changsha, Hunan, PR China.

LIU Yong, College of Sports Science, Changsha Normal University, Changsha, Hunan, PR China.

Contributions

The authors contributed equally to this study. The first author made substantial contributions to the conception and design, or acquisition of data, or analysis and interpretation of data. The second author has been involved in drafting the manuscript or revising it critically for important intellectual content. The third author proofreads and modifies the details of the article.

Corresponding author Correspondence to Wangling.

## Ethics declarations

### Ethics approval and consent to participate

All study procedures, including participant recruitment, informed consent, testing protocols, and safety contingency plans, were reviewed and approved by the Human Research Ethics Committee of East China Normal University (approval no. HR 342-2024).

### Consent for publication

No individual-level data are reported in this study. Not applicable.

